## Extended Data Fig. 1, Extended Data Fig. 2, Table S1 for "Direct observation of autoubiquitination for an integral membrane ubiquitin ligase in ERAD"

Extended Data Fig. 1 Validation of an improved Hrd1 labeling strategy.

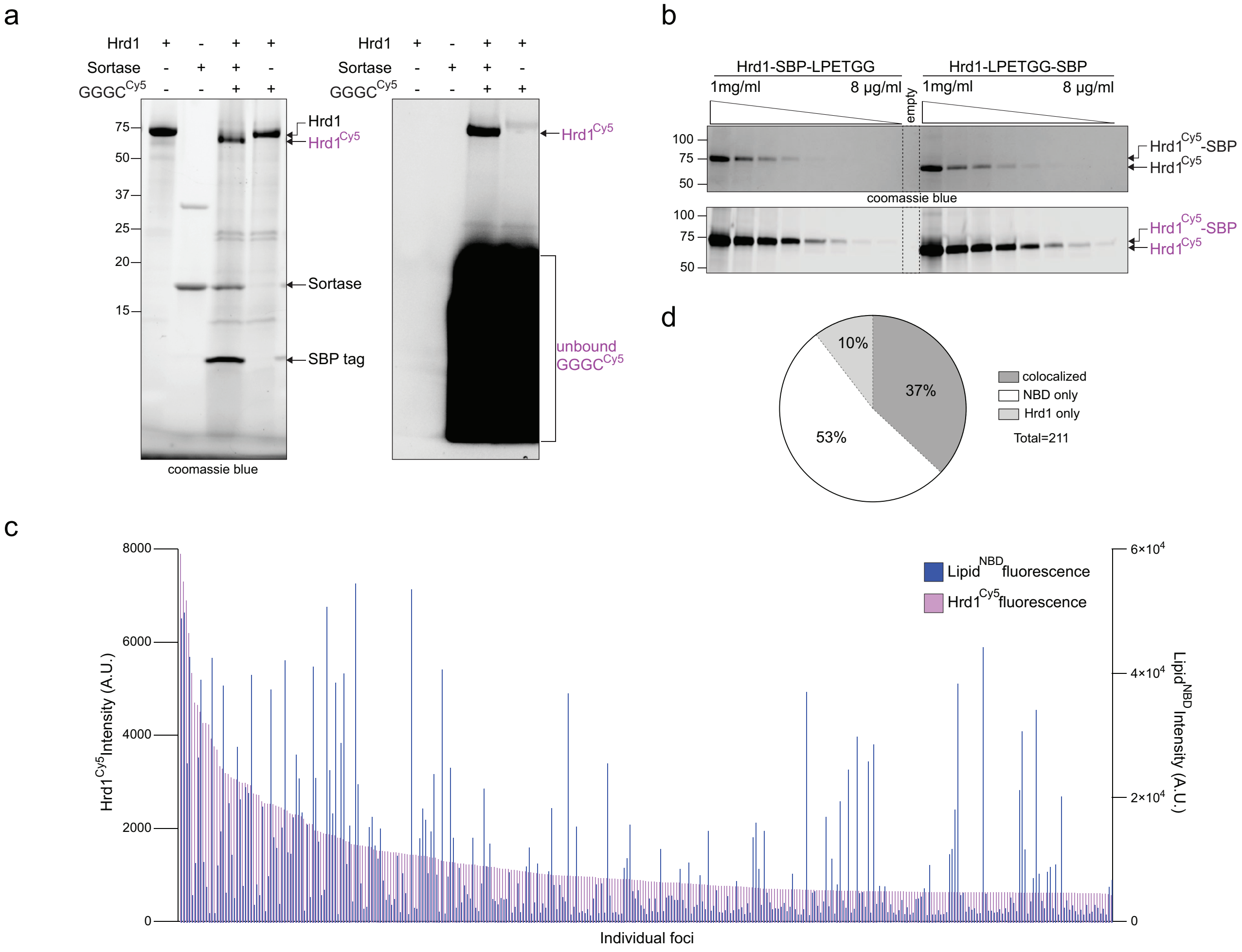

**Supplemental Information**

**Extended Data Fig. 1 Validation of an improved Hrd1 labeling strategy. Related to Figure 1.**

a) Uncropped SDS-PAGE image from Fig. 1a showing additional components of the labeling reaction including sortase A, the cleaved SBP tag, and uncoupled GGGC$^{Cy5}$ peptide.

b) Comparison of the Hrd1 Cy5 labeling efficiency for Hrd1-SBP-LPETGG (previous method) versus Hrd1-LPETGG-SBP (updated method).

c) Colocalization of Hrd1$^{Cy5}$ and lipid$^{NBD}$ in reconstituted proteoliposomes. The three categories correspond to liposomes containing both Hrd1$^{Cy5}$ and lipid$^{NBD}$, liposomes containing only lipid$^{NBD}$ and liposomes containing only Hrd1$^{Cy5}$.

d) Paired fluorescence intensities of Hrd1$^{Cy5}$ and the corresponding lipid$^{NBD}$. Proteoliposome foci were sorted based on Hrd1$^{Cy5}$ intensities.

Extended Data Fig. 2 Additional characterization of the Poisson dilution based counting platform.

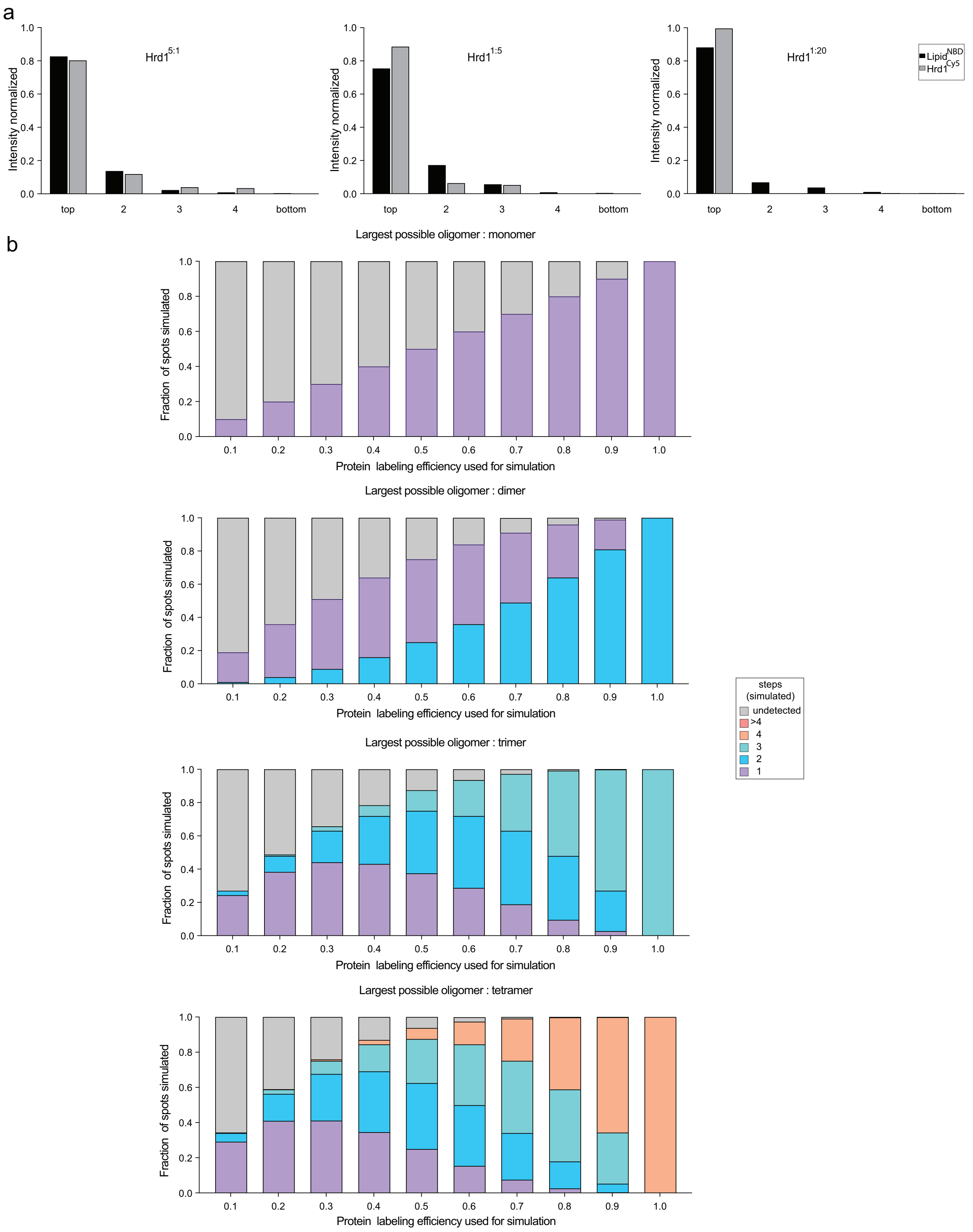

**Extended Data Fig. 2. Additional characterization of the Poisson dilution based counting platform. Related to Figure 2.**

a) $Hrd1^{Cy5}$ proteoliposomes were reconstituted at three different protein:lipid ratios and floated using a glycerol density gradient. The gradient fractions were collected and analyzed using a fluorescence emission based plate reader to track the lipid and Hrd1 migration across the different glycerol layers. Fluorescence values for $Hrd1^{Cy5}$ and $lipid^{NBD}$ were normalized to their respective total intensity from all five fractions for each reconstitution condition.

b) Simulated binomial distributions for monomeric, dimeric, trimeric or tetrameric complex plotted across varying protein labeling efficiencies displayed as simulated step distributions of various-sized complexes. The simulated step distribution in Fig. 2d represents the data corresponding to 90% labeling efficiency across these four separate graphs.

Extended Data Fig. 3 Ubiquitination reaction and correlation plots supporting the single-molecule ubiquitination experiments.

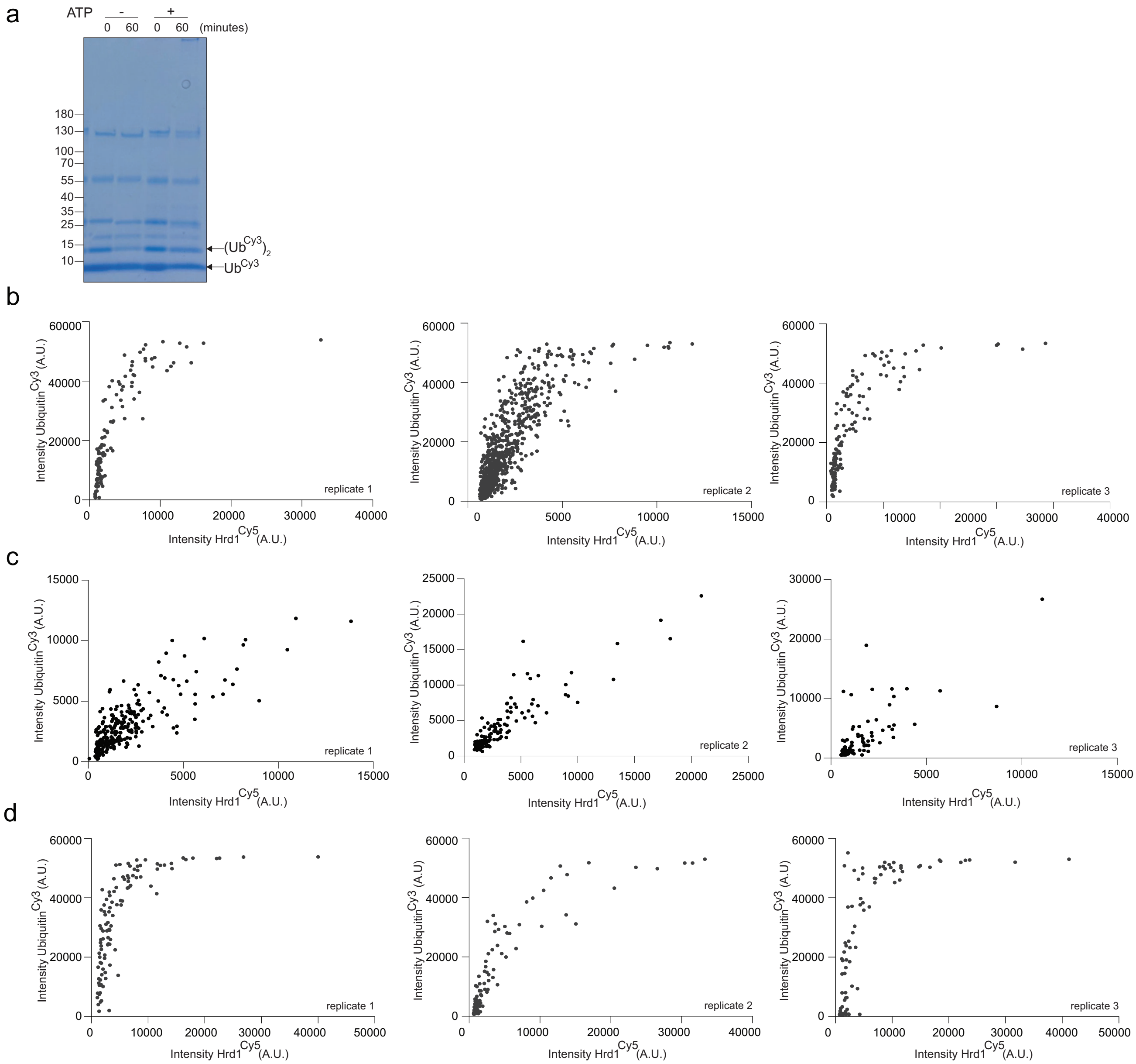

**Extended Data Fig. 3. Ubiquitination reaction and correlation plots supporting the single-molecule ubiquitination experiments.**

a) Coomassie blue stained gel from Fig. 3a.

b) Correlation plots of fluorescence intensities comparing Cy3 (fluorescent label on ubiquitin) and Cy5 (fluorescent label on Hrd1) for ubiquitinated $Hrd1^{20:1}$ proteoliposomes from three independent experiments.

c) Correlation plots of fluorescence intensities comparing Cy3 (fluorescent label on the tandem ubiquitin binding element, TUBE) and Cy5 (fluorescent label on Hrd1) for ubiquitinated $Hrd1^{20:1}$ proteoliposomes from three independent experiments.

d) Correlation plot of fluorescence intensities of Cy3 (fluorescent label on ubiquitin) and Cy5 (fluorescent label on Hrd1) for ubiquitinated $Hrd1^{20:1}$ proteoliposomes diluted with 20 fold excess of empty liposomes from three independent experiments.

**Table S1. Plasmids used in this study**

| Plasmid name | Backbone, features | Reference | Figures |
|---|---|---|---|
| pBMA003 | N terminal $His_{14}$-SUMO, N terminal Cysteine for maleimide labeling | 25 | Figs 3,4 |
| pRSET-6xTR-TUBE | N-terminal His6-T7 tag (N terminal on insert), 6 tandem repeats of trypsin-resistant UBQLN1 UBA domain, all Arg residues in the UBA domain are mutated to Ala, | 46<br><br>Addgene #110313 | Fig 3 |
| pBMA15 | pR426 vector, Gal1 promoter, C terminal Sortase tag, 3 C cleavage site, Streptavidin binding protein tag, Cyc1 terminator | This study | Figs 1,2,3,4 |
| p30 | pET28b vector, Ubc7 with N-terminal $His_6$ tag | 25 | Figs 3,4 |
| p32 | pET28b vector, truncated soluble Cue1 (24-203) N-terminal $His_6$ tag | 25 | Figs 3,4 |
| p37 | Uba1 with N terminal $His_{14}$-SUMO tag | 25 | Figs 3,4 |
